# Evolutionary Decoding of the *Bacillus subtilis* Secretome: Insights from Pan-Genomics and Deep Learning

**DOI:** 10.64898/2025.12.19.695619

**Authors:** Chong Peng, Chengwu Yuan, Linhao Meng, Yuying Chen, Shuang Wen, Fuping Lu, Yihan Liu

## Abstract

*Bacillus subtilis* serves as a crucial host for industrial protein production, where the efficiency and regulation of its secretion system represent a central focus of applied research. Despite substantial genomic diversity among strains, current understanding of signal peptides, which are key elements in the secretion process, remains largely based on single model strains, lacking systematic investigation from a pan-genomic perspective. This study integrates pan-genomics and deep learning to construct a pan-secretome profile of 287 *B. subtilis* strains.

Our analysis reveals an open repertoire of signal peptides in *B. subtilis.* Our results reveal an open pan-signal peptidome in *B. subtilis*. Core signal peptides exhibit high sequence conservation and primarily direct the localization of housekeeping proteins, whereas accessory signal peptides govern the extracellular secretion of environment-responsive proteins, forming a functional “housekeeping-adaptive” dichotomy. We demonstrated for the first time at a broad scale the widespread evolutionary decoupling between signal peptides and their corresponding mature peptides, uncovering a modular evolutionary mechanism in protein evolution. While the deep learning model achieved high accuracy (88.0%) in discriminating core and accessory gene-encoded full-length proteins, its performance was considerably limited when using signal peptide sequences alone (69.7% accuracy), reflecting the information constraints inherent to short sequences.

This study provides a systematic portrayal of the signal peptide evolutionary landscape in *B. subtilis* at the pan-genome scale, advancing our understanding of the evolutionary principles governing bacterial secretion systems and establishing a theoretical foundation for optimizing industrial protein production through rational signal peptide design.

## 1. Introduction

*Bacillus subtilis*, a Gram-positive model bacterium, is recognized as an ideal host for the production of industrial enzymes and high-value compounds due to its non-pathogenic nature, robust protein secretion capacity, and FDA GRAS (Generally Recognized as Safe) status^[1]^. As research into its application potential deepens, scientists have increasingly recognized that phenotypic heterogeneity among different *B. subtilis* strains, such as variations in enzyme yield and environmental adaptability, is closely linked to their genomic diversity. This diversity not only influences industrial performance but also shapes the evolutionary trajectory of their secretory systems ^[2]^.

Advances in high-throughput sequencing have provided new opportunities for deciphering bacterial genomic diversity ^[3]^. Genomic studies have revealed that even within the same bacterial species, genomes can vary significantly among strains inhabiting different environments and hosts ^[4]^. To address this, Tettelin et al. proposed the concept of the “pan-genome,” which partitions the gene repertoire of a species into the core genome (shared by all strains and essential for basic physiological functions) and the accessory genome (found only in a subset of strains and contributing to environmental adaptability) ^[5–7]^. This framework offers a novel perspective for understanding microbial evolution.

Recent pan-genome studies of *B. subtilis* have yielded key insights: Neal et al., through metabolic modeling of 481 strains, revealed that the strains could be classified into five functional groups according to their metabolic characteristics, elucidating the mechanisms by which they adapt to specific environments ^[8]^. Kim et al. demonstrated via comparative genomics of 238 *Bacillus* strains that core genes are functionally conserved, while accessory genes drive strain-level diversity ^[9]^. Wu Hao et al. proposed strain screening optimization strategies, significantly enhancing the reliability of pan-genome analyses^[10]^. These findings not only deepen our understanding of the evolutionary mechanisms of *B. subtilis* but also provide theoretical support for targeted strain breeding and ecological function exploration.

The industrial value of *B. subtilis* hinges largely on its highly efficient protein secretion system. Through the classical Sec and Tat (twin-arginine translocation) pathways, the bacterium directly transports target proteins like amylases and proteases to the extracellular milieu, substantially simplifying downstream processing and reducing costs ^[11–14]^. The signal peptide (SP), a critical component guiding nascent proteins to the membrane or for export, not only determines secretion efficiency but is also deeply involved in environmental adaptation and host interactions ^[15]^. However, current research faces notable limitations. Most conclusions are still drawn from single model strains, failing to adequately account for the impact of strain diversity on signal peptide function. Moreover, systematic insights into the evolutionary dynamics of signal peptides are lacking, particularly regarding the balance between the conservation of core signal peptides and the plasticity of accessory ones. Although pan-genomes has been successfully applied to elucidate metabolic diversification and environmental adaptation in *Bacillus*, its potential for advancing secretion system research remains largely underexplored.

Currently, pan-genome analyses have not been deeply integrated with secretome studies, leaving patterns of signal peptide conservation at the species level uncharacterized. Concurrently, emerging deep learning techniques, despite their groundbreaking potential for gene classification and prediction, have never been applied to pan-genome analysis or core gene prediction.

This study bridges three critical domains: i) constructing a pan-secretome profile based on 287 *B. subtilis* strains; ii) systematically deciphering the conservation characteristics of signal peptides from core and accessory genes; and iii) developing deep learning models to distinguish between core and accessory genes, as well as conserved versus adaptive signal peptides. By revealing the “function-plasticity” balance mechanism of signal peptides under evolutionary pressure, this work provides a theoretical basis for strain optimization in metabolic engineering and industrial applications, thereby contributing to the development of more efficient protein expression and secretion systems.

## 2. Materials and methods

### 2.1 Genomic data acquisition and annotation

All complete *Bacillus subtilis* genomes were downloaded from NCBI (as of February 2024)[^16]^。To ensure data quality, only fully assembled strains were retained, and those with an N-base content exceeding 1% were filtered out using a Python script, resulting in 287 high-quality genomes. Gene prediction and functional annotation were performed using Prokka [^17].^

### 2.2 Pan-genome construction and signal peptide prediction

The pan-genome was constructed from Prokka-generated files using Roary ^[18]^ (BLASTP identity threshold: 95%) to identify core and accessory genes. Signal peptides were predicted for all encoded proteins using SignalP 6.0 ^[19]^.

### 2.3 Signal peptide characterization and functional analysis

Sequence similarity matrices for core and accessory genes were computed using Clustal Omega ^[20]^, with mean similarity values compared via t-test to assess conservation differences. Functional enrichment analysis (GO and KEGG) for signal peptide-containing genes was conducted using the DAVID database^[21]^.

### 2.4 Protein localization and phylogenetic visualization

Subcellular localization was predicted with PSORTb^[22]^ and visualized as proportion charts ^[23]^. A genome-wide phylogenetic tree was built using OrthoFinder ^[24]^ and visualized with iTOL ^[25]^, overlayed with signal peptide distribution heatmaps to illustrate evolutionary-functional relationships.

### 2.5 Deep learning-based protein classification

To evaluate the predictive capability of deep learning for identifying core proteins and signal peptides in core proteins, we constructed three distinct datasets for model training. Based on pan-genome annotations derived from 287 strains, we identified 1,884 core genes and 5,936 unique genes. Since core genes are present in all strains, they collectively yielded approximately 541,000 protein sequences. To eliminate redundancy and obtain representative sequences, we applied CD-HIT ^[26]^ with a sequence similarity threshold of 0.9 for clustering, ultimately retaining 1,884 core protein sequences and 5,936 unique protein sequences to form the Core/Unique Protein dataset.

Subsequently, we filtered this dataset to retain only proteins containing signal peptides, resulting in 101 core proteins and 227 unique proteins which contains signal peitides. From these, we constructed two additional signal peptide-related datasets: one comprising the signal peptide sequences alone, and the other containing the full-length amino acid sequences of these proteins. Each dataset was partitioned into training and test sets using an 8:2 ratio, while maintaining balanced distribution between the two protein categories.

Protein language model ESM-2 ^[27]^ was used to generate embedding vectors for sequences. A Support Vector Machine (SVM) with RBF kernel was employed for classification, with hyperparameters optimized via grid search. Model performance was evaluated using accuracy (ACC), sensitivity (SN), specificity (SP), Matthews correlation coefficient (MCC), and area under the ROC curve (AUC). Feature embeddings were visualized in two-dimensional space using UMAP ^[28]^.

## 3. Results

### 3.1 Analysis of signal peptide diversity in the pan-genome framework of *Bacillus subtilis*

To elucidate the diversity and evolutionary conservation of genes and their signal peptides in *Bacillus subtilis*, we performed a comprehensive pan-genome analysis of 287 strains, with particular emphasis on signal peptide sequences.

The gene family accumulation curve demonstrated an open pan-genome structure for *B. subtilis*, where the total number of gene families increased continuously without reaching a plateau as more genomes were added (Figure 1A). This pattern indicates that newly incorporated strains consistently contribute non-conserved genes, reflecting the organism’s active gene acquisition capability and substantial genetic plasticity. In contrast, the core genome size (red curve) eventually stabilized, suggesting a limited set of conserved gene families maintained across all strains.

**Figure 1.**
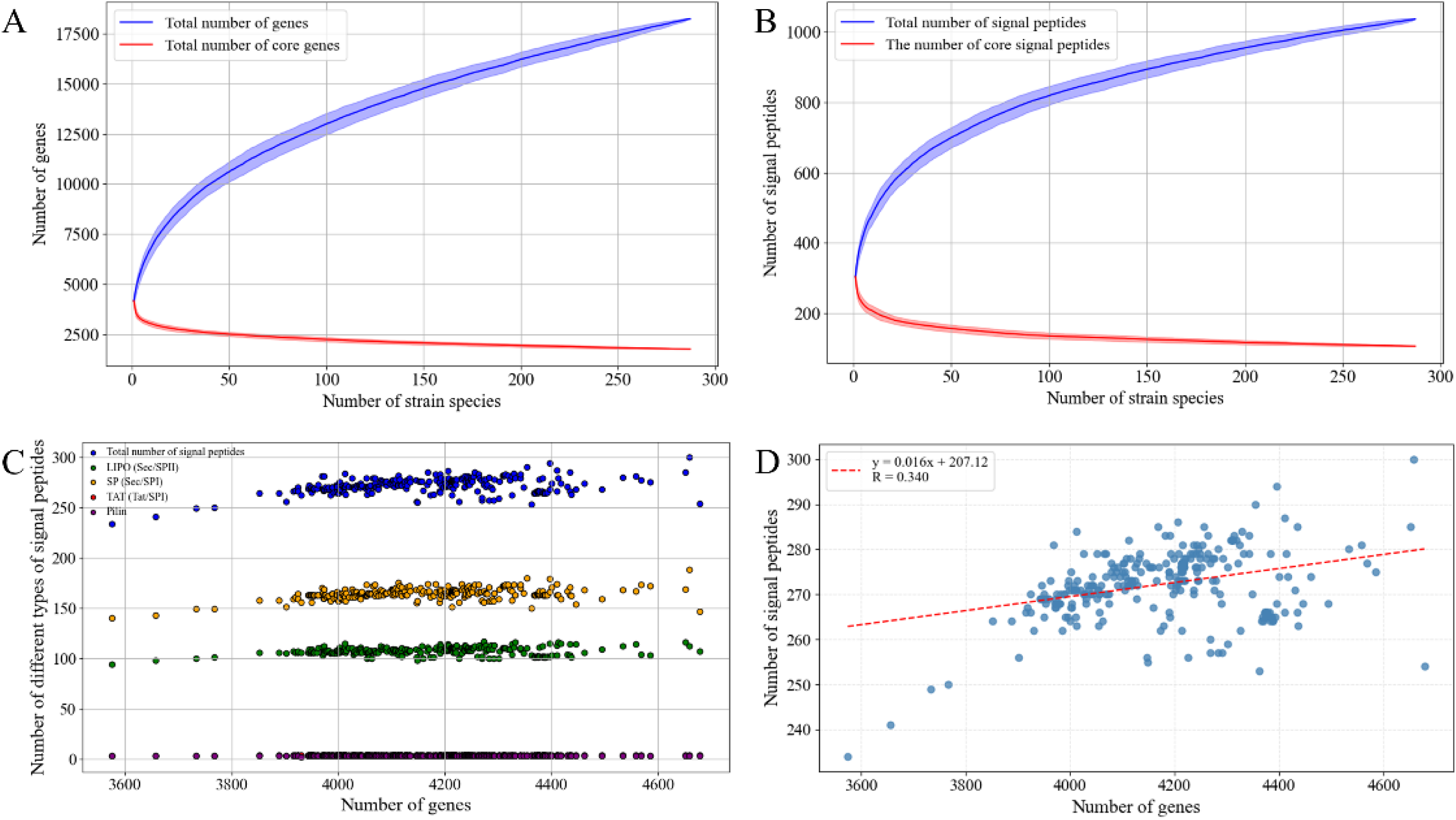
Pan-genome and signal peptide analysis across 287 *Bacillus subtilis* strains. (A) Gene family accumulation curves, displaying the trends in total gene count (blue line) and core gene count (red line) as the number of strains increases. (B) Signal peptide accumulation curves, illustrating the trends in total signal peptides (blue line) and core signal peptides (red line) with increasing strain numbers. (C) Scatter plot showing the relationship between total gene count and the abundance of different signal peptide types, including Sec/SPII prokaryotic lipoprotein signal peptides (LIPO, green), Sec/SPI standard secretory signal peptides (SP, orange), Tat/SPI twin-arginine signal peptides (TAT, red), and Sec/SPIII Pilin-type signal peptides (purple). Blue dots represent the total signal peptide count. (D) Scatter plot depicting the correlation between gene number and signal peptide number across different strains.

**Table 1.** Classification performance of the deep learning model on three datasets.

| Dataset | Data size | ACC | SN | SP | MCC | AUC |
| --- | --- | --- | --- | --- | --- | --- |
| Core/Unique Protein | 1,884 core<br>/5,936 unique | 0.880 | 0.928 | 0.729 | 0.667 | 0.928 |
| SP sequences from Core/Unique Proteins | 101 core/ 227<br>unique | 0.697 | 0.804 | 0.450 | 0.262 | 0.689 |
| Full-length amino acid sequences of Core/Unique Proteins containing SPs | 101 core/ 227<br>unique | 0.773 | 0.717 | 0.900 | 0.815 | 0.860 |

A closely analogous trend was observed in the analysis of signal peptide abundance (Figure 1B). The total number of distinct signal peptides also increased steadily without reaching saturation as more strains were added, suggesting continuous introduction of non-conserved signal peptides by new isolates. The number of core signal peptides stabilized at approximately 200 after the inclusion of around 100 strains, demonstrating strong conservation of the fundamental functional components of the secretion system. Notably, accessory (non-core) signal peptides accumulated at a significantly faster rate than core ones, highlighting the rapid evolution of environment-specific secretion functions that underpin strain differentiation. This core-accessory dichotomy reveals the evolutionary logic of bacterial secretion systems: conserved core elements sustain fundamental secretion capacity, while accessory components enable niche-specific adaptation.

Analysis of the relationship between the number of signal peptides and genome size across *B. subtilis* strains revealed a general positive correlation (Figures 1C and 1D). As the number of genes increased, the total repertoire of signal peptides expanded correspondingly, suggesting that genomic expansion is typically accompanied by enhanced secretory demands. However, beyond approximately 4,200 genes, the rate of signal peptide accumulation decelerated, implying potential functional or regulatory constraints.

Among the four canonical signal peptide types, Sec/SPI (standard secretory) was the most abundant and exhibited sustained growth with increasing gene numbers, reflecting its role as the primary secretory pathway. Sec/SPII (lipoprotein-type) was the second most prevalent. In contrast, Tat/SPI (twin-arginine) and Sec/SPIII (Pilin-type) signals were fewer in number and sporadically distributed. The latter was particularly rare, present in only single-digit quantities or entirely absent in most samples, indicating highly specialized functions with limited applicability.

In summary, during genomic expansion in *B. subtilis*, the signal peptide system demonstrates both the universal enhancement of fundamental secretory pathways and the restricted distribution of specialized routes. This pattern reflects an evolutionary strategy that maintains core secretory functions while dynamically utilizing accessory elements to adapt to diverse ecological niches.

### 3.2 Characterization of core signal peptide sequences and functional profiles

To investigate the distinctions between signal peptides from core and accessory genes in *B. subtilis*, we systematically compared their sequence characteristics and functional distributions based on pan-genome data.

Sequence conservation analysis revealed that core signal peptides exhibit greater stability. The amino acid mutation rate of core signal peptides was significantly lower than that of accessory signal peptides (Figure 2A), consistent with the overall evolutionary conservation of core genes. This indicates that sequences essential for maintaining basic secretory function are subject to stringent constraints. Furthermore, comparative analysis of mutation patterns between signal peptides and their corresponding mature peptides showed that signal peptides display a mutation profile characterized by a lower median but higher dispersion. This pattern suggests strong overall selective pressure to maintain function, with some tolerance for variation under certain conditions or proteins. In contrast, mature peptides exhibited a higher median mutation rate with lower dispersion, reflecting a more balanced evolutionary model between conserved domains and variable regions as the functional core of the protein. These findings indicate that in engineering signal peptides for protein production, critical residues must be rigorously conserved, while non-essential regions offer opportunities for rational optimization.

**Figure 2.**
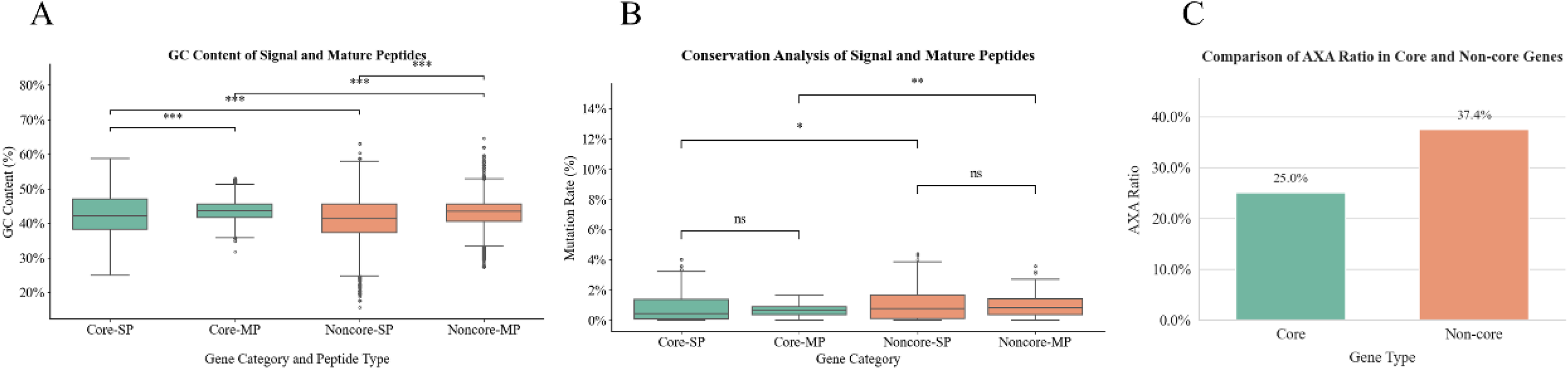
Sequence characteristics of signal peptides derived from core proteins. (A) Box plots comparing the sequence conservation of signal peptides from core and accessory proteins, along with their corresponding mature peptides. (B) GC content distribution in the DNA sequences encoding core and accessory signal peptides and their mature peptides. (C) Proportion of core and accessory signal peptides terminating with the “AXA” motif.

GC content analysis highlighted functional specialization between signal peptides and mature peptides. Significant differences in GC content were observed among the four sequence groups: core signal peptides, core mature peptides, accessory signal peptides, and accessory mature peptides (Figure 2B). Regardless of gene conservation status, signal peptides systematically exhibited lower GC content than their corresponding mature peptides. This pattern stems from the requirement for the hydrophobic core region (h-region) of signal peptides to be enriched with non-polar amino acids encoded predominantly by AT-rich codons. Consequently, low GC content represents an inherent requirement for membrane targeting function and may additionally optimize mRNA structure to facilitate recognition by signal peptidase. In contrast, the relatively higher and stable GC content of mature peptides aligns more closely with the host genomic background, thereby supporting translation efficiency and long-term functional stability.

Analysis of the signal peptidase cleavage site further elucidates functional differentiation. Previous studies have indicated that the C-terminal AXA motif of signal peptides correlates with enhanced secretion efficiency of heterologous proteins ^[29]^.

Our quantification of this tripeptide’s prevalence revealed a significantly lower proportion of AXA-containing core signal peptides compared to their accessory counterparts (Figure 2C). We hypothesize that the AXA motif primarily optimizes extracellular protein secretion. Since core proteins predominantly function at cellular membranes or within the periplasmic space, their reduced demand for highly efficient extracellular export explains their lower utilization of this extracellular secretion-biased cleavage motif. Conversely, accessory proteins frequently employ AXA as a “universal efficiency module” to facilitate rapid extracellular export, enabling adaptation to complex environmental conditions and competitive pressures. This finding, at the level of signal peptide sequence characteristics, validates the functional differentiation in secretion destinations and evolutionary strategies between core and accessory proteins.

Functional enrichment and localization analyses revealed fundamentally distinct biological roles for core and accessory signal peptides. GO enrichment analysis demonstrated that core proteins containing signal peptides were predominantly associated with essential cellular maintenance processes, including sporulation, cell wall organization, and peptidoglycan turnover (Figure 3A). Their molecular functions involved serine-type endopeptidase activity, with cellular localization primarily concentrated at the plasma membrane and periplasmic space (Figure 3B). These characteristics identify them as housekeeping elements crucial for maintaining cellular structural integrity and basic physiological activities.

**Figure 3.**
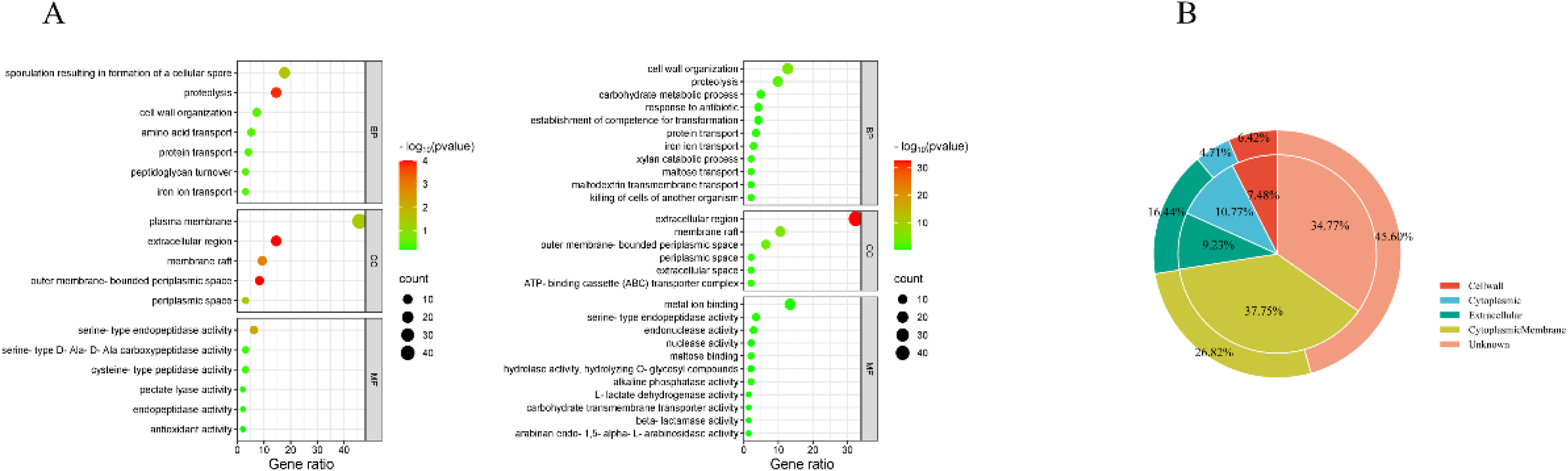
Functional divergence and subcellular localization differences between core and accessory proteins in Bacillus subtilis. (A) GO enrichment analysis of core proteins (left panel) and accessory proteins (right panel). The x-axis represents the gene ratio, while the y-axis shows significantly enriched GO terms. Dot size corresponds to the number of enriched genes in each term, and color intensity indicates enrichment significance. (B) Proportional distribution of subcellular localization for core and accessory proteins. The inner ring displays core proteins, and the outer ring represents accessory pproteins, shown in a pie chart format.

In contrast, accessory proteins containing signal peptides were predominantly enriched in environmental adaptation processes such as antibiotic response, xylan catabolism, and maltose transport. They showed significant association with the “killing of cells of another organism” function, while their molecular functions included pectate lyase and antioxidant activities. Their localization preferentially targeted the extracellular region and membrane rafts, indicating that accessory signal peptides serve as adaptive elements mediating environmental competition and nutrient acquisition, thereby facilitating niche expansion.

In summary, core and accessory proteins containing signal peptides in *Bacillus subtilis* demonstrate clear differentiation in both sequence evolution and biological function: core signal peptides exhibit sequence conservation and inward-oriented functions dedicated to maintaining fundamental cellular structure and homeostasis, whereas accessory signal peptides display sequence plasticity and outward-oriented functions driving environmental adaptation and interspecies competition.

### 3.3 Evolutionary decoupling between signal peptides and mature peptides

To investigate the evolutionary relationship between signal peptides and mature peptides, we analyzed sequence variations in three core genes (SacB, YwnA, and PelC) across 287 *Bacillus subtilis* strains. By mapping sequence identity of signal peptides and their corresponding mature peptides onto a phylogenetic tree (Figure 4A), we observed a remarkable phenomenon: strains with identical signal peptide sequences frequently exhibited substantial divergence in their corresponding mature peptides. For instance, while the SacB signal peptide was completely conserved across all strains (uniform blue band), its mature peptide displayed multiple distinct variants (multi-colored band). This pattern, consistently observed across all three genes examined, indicates widespread evolutionary decoupling between signal peptides and mature peptides.

**Figure 4.**
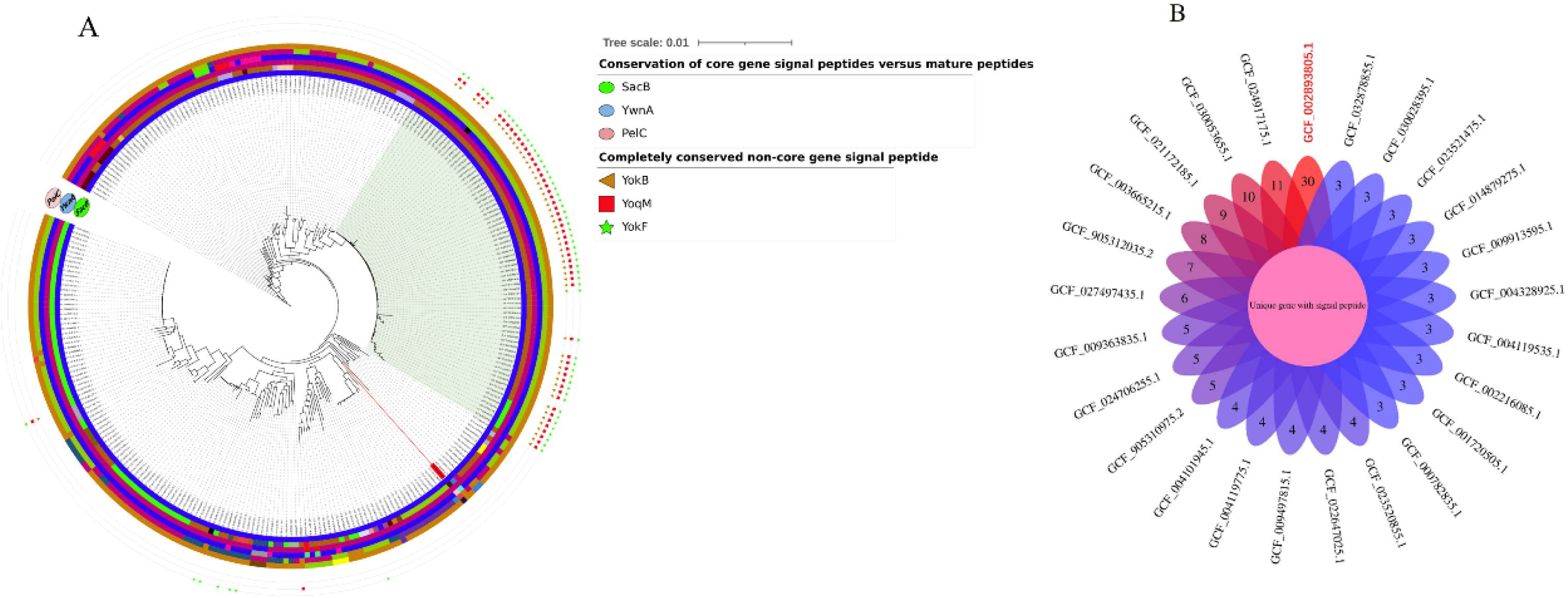
Phylogenetic analysis of *Bacillus subtilis* with signal peptide conservation and specificity profiles. (A) Phylogenetic tree of *B. subtilis* strains. The concentric rings from inner to outer represent signal peptides and mature peptides of SacB, YwnA, and PelC, respectively. Identical signal peptide sequences across strains are marked by the same color in corresponding bands, following the same scheme for mature peptides. Color shifts within bands indicate sequence variations in signal peptides/mature peptides compared to adjacent strains. The outermost symbols denote perfectly conserved accessory gene signal peptides, with their corresponding strains showing clustering tendencies on the phylogenetic tree. (B) Petal chart displaying the number of unique genes containing signal peptides across different strains.

This decoupling likely stems from their distinct selective constraints. Signal peptides, serving as targeting sequences for protein secretion, evolve under stringent constraints imposed by the conserved physicochemical rules of the secretory machinery, maintaining high sequence conservation among closely related strains. In contrast, mature peptides which house the functional domains evolve primarily under ecological pressures, driving sequence diversification across strains as they adapt to different niches. This modular evolutionary mechanism enables bacteria to rapidly generate functional variants through signal peptide swapping or mature peptide recombination, bypassing the need for whole-gene rearrangements and thereby significantly accelerating evolutionary innovation.

Our analysis also identified accessory genes (YokB, YoqM, and YokF) whose signal peptides maintained 100% sequence identity across all strains carrying them. Phylogenetic analysis revealed that strains containing these perfectly conserved signal peptides formed distinct clusters on the evolutionary tree (outer markers in Figure 4A), suggesting their dissemination through horizontal gene transfer (HGT). Particularly noteworthy is the prophage-derived YokB gene, whose perfectly conserved signal peptide serves as a molecular marker for HGT events, providing reliable evidence for reconstructing strain evolutionary history.

Strain GCF_002893805.1 deserves special attention, exhibiting a significantly elongated branch on the phylogenetic tree (Figure 4A), indicating a unique evolutionary trajectory. Isolated from the high-salt, protein-rich environment of Chinese food lobster sauce, this strain likely underwent rapid adaptive evolution within this relatively confined niche. Genomic analysis revealed that it encodes 30 unique signal peptide-containing genes, substantially exceeding counts in other strains (Figure 4B). These genes encode various hydrolytic enzymes (including Endo-1,4-beta-xylanase A and Endoglucanase), salinity stress proteins (K(+)/H(+) antiporter modulator KhtS), and biofilm-associated proteins, all potentially contributing to its adaptation within this distinctive fermentation environment.

Collectively, these findings illuminate the pivotal role of secretory proteins in microbial niche specialization from an evolutionary perspective. The evolutionary patterns of signal peptides, as key regulatory components of the secretion system, directly reflect microbial adaptive strategies to environmental interactions. The abundance of unique signal peptide genes in specialized strains not only demonstrates environment-driven adaptive evolution but also provides valuable insights into how microbes rapidly acquire novel functions through horizontal gene transfer and positive selection.

### 3.4 Deep learning-based classification of core and accessory genes

To overcome the limitations of conventional sequence analysis methods in identifying rapidly evolving genes, we implemented a deep learning framework to extract deep functional semantic features from sequences for accurate discrimination between core and accessory genes.

Using pan-genome data from 287 *Bacillus subtilis* strains, we constructed a dataset comprising protein sequences from 1,884 core genes and 5,936 unique genes as positive and negative samples, respectively. Sequence features were extracted using the ESM-2 protein language model, and a support vector machine (SVM) classifier was employed for the classification task. The model achieved 88.0% accuracy with an AUC of 0.928 in distinguishing core from unique gene-encoded proteins on an independent test set, demonstrating significant separability between these two categories in the feature space.

To visualize the distribution patterns of these features, we applied UMAP for dimensionality reduction. While the original ESM-2 embeddings showed a partial separation trend, substantial overlap remained between the two sample classes (Figure 5A). In contrast, the feature representations learned through SVM training exhibited much clearer clustering structures (Figure 5B), indicating that supervised learning effectively enhances feature discriminability.

**Figure 5.**
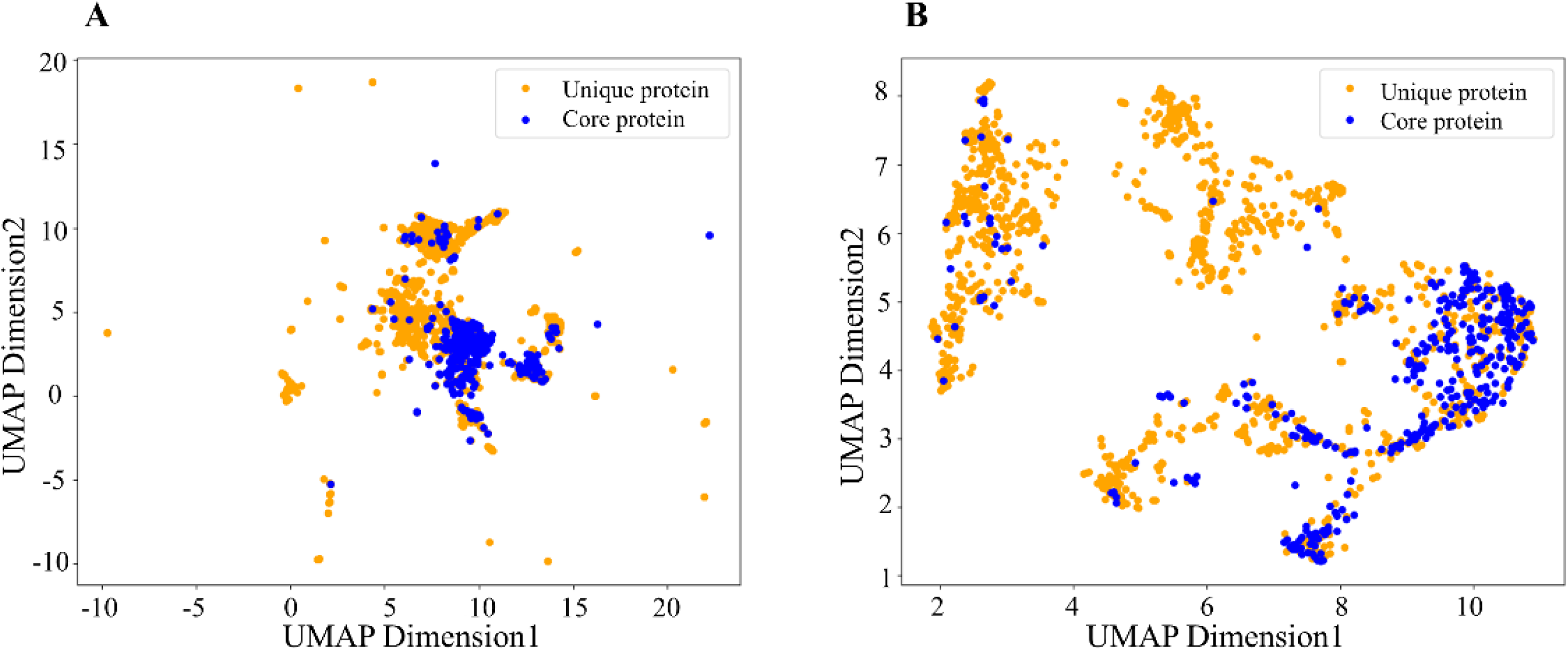
UMAP visualization of ESM-2-derived protein sequence embeddings before (A) and after training (B).

We subsequently extended this analytical framework to both full protein sequences and isolated signal peptide sequences from core and accessory proteins containing signal peptides. Using datasets of 152 core and 194 accessory proteins (and their corresponding signal peptides), classification models achieved accuracies of 77.3% for full protein sequences and 69.7% for signal peptide sequences alone. This performance limitation likely stems from the abbreviated length of signal peptides, constrained dataset size, and inherent structural and functional similarities between core and accessory signal peptides, collectively challenging the effectiveness of sequence-based machine learning approaches for this specific classification task.

This study demonstrates that deep learning can effectively capture deep feature differences between core and accessory gene-encoded proteins, while highlighting remaining challenges in classification at the signal peptide level, thereby providing direction for future research endeavors.

## 4. Discussion

This study establishes an integrated framework combining pan-genomics and deep learning to systematically investigate signal peptide evolution in *Bacillus subtilis*. Our findings reveal a fundamental “core-accessory” dichotomy in signal peptide evolution, demonstrating how bacteria employ differentiated secretion strategies to balance genetic stability with environmental adaptability.

We observed remarkable consistency between signal peptide evolution and the pan-genome architecture. Core signal peptides demonstrate significant sequence conservation and functional specialization in essential cellular processes including sporulation and cell wall organization, functioning as housekeeping elements that maintain physiological homeostasis. Conversely, accessory signal peptides are predominantly associated with environmental adaptation mechanisms such as antibiotic response and substrate degradation, serving as modular, adaptive components that enable niche expansion. This functional dichotomy indicates that *B. subtilis* ensures survival baseline by maintaining stable core secretion mechanisms while utilizing accessory component diversity to achieve niche specialization.

Notably, we identified widespread evolutionary decoupling between signal peptides and their corresponding mature peptides. In numerous core proteins, signal peptides remain highly conserved while mature peptides exhibited substantial variability. This pattern indicates differential selective pressures on distinct protein functional modules: signal peptides evolve under stringent physicochemical constraints of the secretory machinery, functioning essentially as conserved “passports” for membrane translocation, whereas mature peptides experience direct environmental selection as functional entities. This modular evolutionary strategy enables rapid functional innovation through horizontal gene transfer, representing a crucial mechanism for bacterial adaptation.

The deep learning approach successfully distinguished between core and accessory gene-encoded proteins with high accuracy (88.0%), indicating fundamental differences in their underlying feature representations. However, classification performance decreased substantially when using only signal peptide sequences (69.7% accuracy), likely reflecting the limited information content of these short sequences and potential functional similarities between core and accessory signal peptides.

While this study provides significant insights, certain limitations should be acknowledged, including the potential for expanding strain diversity and the need for experimental validation of conclusions. Looking forward, deeper understanding of signal peptide evolutionary logic will directly benefit synthetic biology applications, such as rational design of high-efficiency signal peptides to optimize secretion capacity in industrial strains. Furthermore, perfectly conserved signal peptides in accessory genes can serve as molecular signatures for horizontal gene transfer, providing novel tools for precise reconstruction of strain evolutionary history.

